# Orientation Dependence of R_2_’ in the White Matter: Digital Characterization, Modelling and Implications for Studying Brain Physiology

**DOI:** 10.64898/2026.06.02.729601

**Authors:** Nayana Menon, Xiaole Z. Zhong, Yutong Lydia Sun, J. Jean Chen

**Author notes:** Corresponding author: Nayana Menon.

## Abstract

**Purpose:** R_2_*, the transverse relaxation rate, reflects local magnetic field inhomogeneities from susceptibility differences with R2’, the reversible component sensitive to blood oxygenation. Orientation dependence of R_2_ and R_2_* in white matter (WM) are attributed primarily to myelin, with vascular contributions to R2’ uncharacterized. This study examined WM R_2_’ orientation dependence, evaluated existing models, and developed an improved model combining myelin and blood.

**Methods:** Simulations used BOLDswimsuite with 2D WM voxels (5,000 fibres). Spin-echo (TE = 70ms) and gradient-echo (TE = 35ms) signals were simulated across 30 fibre orientations (0°–90°). R_2_′ was calculated as R_2_* − R_2_. Oxygenation, cerebral blood volume (CBV), vessel size, and vessel geometry were varied. Four published models and a novel Myelin-Blood model were fitted to R_2_′ data and compared using R^2^ and RMSE.

**Results:** R_2_ and R_2_* showed strong orientation dependence. Parallel and mixed vessel geometries produced greater R_2_′ amplitude and orientation dependence than random geometries; decreasing oxygenation and increasing CBV amplified orientation effects. Vessel size altered peak locations. Existing vascular models performed poorly, and the Empirical Myelin Model erred near the magic angle. The Myelin-Blood model provided near-perfect fits (mean R^2^ = 0.999, RMSE = 0.007 Hz), reducing RMSE by ~74%.

**Discussion:** WM R_2_′ cannot be explained by vascular or myelin effects alone. Myelin is the primary determinant of orientation dependence, but vascular contributions were evident near the magic angle and low oxygenation. The Myelin-Blood model improves WM R_2_’ characterisation and may reduce orientation-dependent bias and improve interpretation of WM BOLD fMRI signals.

## Introduction

Blood-oxygenation level-dependent (BOLD) functional MRI (fMRI) is a well-established technique for mapping brain activity non-invasively ^1,2^. BOLD fMRI detects changes in neural activity through changes in the T_2_*-weighted signal intensity, which reflect changes in R_2_*. R_2_* can in turn be expressed as R_2_* = R_2_ + R_2_’ ^3^; accurate interpretation of BOLD signals therefore requires understanding how R_2_’ reflects the underlying physiology. In grey matter (GM), the dominant susceptibility source is deoxyhemoglobin ^1,2^, and the frequency shift induced by a vessel depends on its orientation relative to the main magnetic field B_0_ ^1,4^. In a voxel containing a number of randomly oriented vessels, these orientation-dependent shifts produce a net phase dispersion that can be approximated as a vessel orientation-independent R_2_’ ^1,3^, and in the GM, the overall orientation is assumed to be non-directional.

WM differs from GM in that it contains densely packed, highly directional myelinated axon bundles, and the myelin sheath has diamagnetic susceptibility ^5–8^. Consistent with this, several studies established that R_2_* in WM varies significantly with fibre orientation ^6,9–11^, and subsequent work at 7T modelled R_2_* as a combination of sin(2θ) and sin(4θ) terms, with the sin(2θ) term attributed to isotropic bulk susceptibility of the myelin sheath and the sin(4θ) term to susceptibility anisotropy, whereby the effective susceptibility (net magnetic response of the sheath) varies with fibre orientation ^5,12,13^. The orientation dependence of R_2_ has also been extensively documented in the literature, with one noted characteristic being the magic angle effect. R_2_ exhibits a characteristic minimum near the “magic angle” (θ = 54.7°) and a maximum at perpendicular orientations, a pattern consistent with both susceptibility based diffusion through mesoscopic field gradients and residual dipolar coupling mechanisms, where rotationally restricted water molecules aligned along axonal microstructure retain a non-averaged intramolecular dipolar interaction that varies with fibre orientation ^14–16^.

While the orientation dependence of R_2_ and R_2_* in WM has been characterized in this way, the effects of blood on orientation dependence in the WM have not been systematically examined. This is consequential for WM fMRI. Although the WM BOLD signal has historically been deemed undetectable due to the slower, lower amplitude hemodynamic responses relative to GM ^17–20^, and although myelin orientation already drives strong orientation dependence in relaxation rates, the recent growth in fMRI calls for a renewed understanding of the vascular contribution to this R_2_’ orientation dependence, and how existing models can be used to correct it.

An additional complexity is that WM blood vessels do not entirely run parallel to the fibres. Anatomical studies have established that approximately 50% of WM vessels run parallel to fibre tracts, with the remaining vessels randomly oriented in the tissue ^21–23^. Also, while WM is considerably less vascularized than GM, with cerebral blood volume (CBV) typically estimated at 2% ^24,25^, large penetrating vessels from the pial surface have been shown to extend into the WM^26^, which could potentially contribute a macrovessel component alongside capillaries in the WM. Moreover, vessel radii in WM have not been systematically characterized histologically, unlike axons and axon radii ^27,28^, leaving the size distribution of WM vessels relatively unknown.

In this work, we use Monte Carlo simulations ^1,5,13,29,30^ to systematically characterize the orientation dependence of R_2_’ in WM while varying the blood oxygenation, CBV, vessel diameter, and axon spacing, and to evaluate how well existing empirical and biophysical models model the simulated data. Monte Carlo simulations account for the diffusion of water protons through spatially varying magnetic fields ^31^, and have been widely used as a way to predict pseudo R_2_ and R_2_’ behaviour. Four published R_2_’ models were compared, including three static dephasing vessel models ^3^ and an empirical myelin model ^12^. In addition, we propose a novel R_2_’ model that incorporates both myelin and vascular compartments and evaluate its performance relative to existing models. By characterizing the vascular contribution and orientation dependence of R_2_’ in WM and evaluating the performance of existing models, this work addresses a gap that has direct consequences for the interpretation of the WM BOLD signal.

## Methods

### Generation of simulated ground truth

Simulations were performed using BOLDsωimsuite^32^, an open-source Python package for forward modelling of BOLD contrast mechanisms (https://github.com/jacobchausse/BOLDswimsuite). The BOLDsωimsuite toolbox was customized to enable each of our WM simulations.

Fibre geometry in an isotropic 2D voxel containing 5000 fibres was generated using the AxonPacking^33^ package where each fibre is represented by a circle (Fig. 1b). For all simulations other than those to test the effect of the vessel diameter, randomly selected circles in Fig. 1b were assigned to the “vessel” class to achieve the CBV necessary.

**Figure 1.**
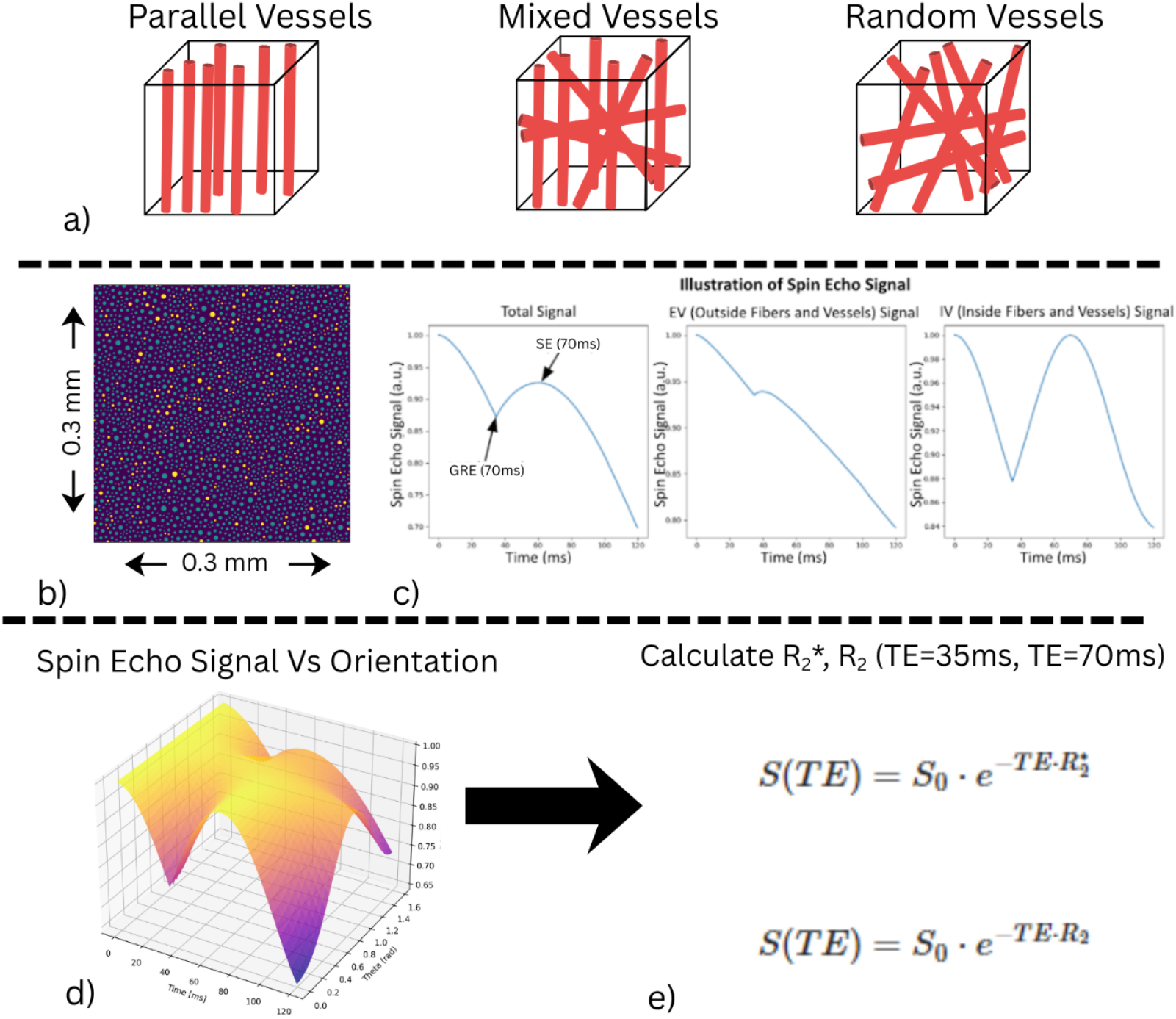
Overview of the simulation methodology. a) Shown here is an example of the vessel orientation structure, shown for a 3D voxel example. b) shows a sample 2D voxel generated in AxonPacking and rendered in BOLDsωimsuite, with a size of 0.3 x 0.3mm. Yellow circles represent blood vessels and green represent myelinated axons, c) Illustrates the simulated total, IV and EV signals from BOLDsωimsuite. d) is a 3D plot of the simulation output which shows the simulated signal across time and voxel orientation relative to the main magnetic field B_0_, e) shows the formulas used to calculate the R_2_* and R_2_, with the signal sampled at 35 and 70ms, respectively.

The simulations encompassed systematic variations in fibre orientation relative to B_0_ (θ = 0° to π/2 in steps of π/64, 30 angles total),blood oxygenation (Y), CBV, vessel diameter and axon spacing, with vessel orientation distribution configurations including parallel, mixed (50% parallel, 50% random), and entirely random vessel orientations (Fig 1a). Default parameter values for each factor are stated in the corresponding subsections below. Haematocrit was fixed at Hct = 0.40 throughout ^34^. The following variables are altered.

### Blood oxygenation

Blood oxygenation (Y) was varied from 0.6 to 0.9 in steps of 0.1.

Oxygenation was reflected in the venous susceptibility (Δχ), ^34^:

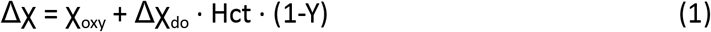

where χ_oxy_ is the susceptibility of oxygenated blood set to −0.008 ppm cgs ^34^ and Δχ_do_ is the susceptibility difference between oxygenated and deoxygenated blood set to 0.27 ppm cgs^34^. Code was added to the BOLDsωimsuite package to accommodate this change.

### CBV

The CBV was varied from 0.01 to 0.03 in steps of 0.01. This range is consistent with previously derived estimates of white matter CBV, which has been shown to be lower than grey matter and typically falls between 1% and 3% ^24,25^. The default value was CBV = 0.02. No modifications were required for this parameter variation.

### Vascular diameter

For the simulations of the vessel diameter effects, axon diameter from the AxonPacking voxel (generated using 0 μm axon spacing) were partitioned into three equal-width bins: small (0.80-3.11 μm), medium (3.11-5.42 μm) and large (5.42-7.72 μm). The maxima and minima of this range are derived from AxonPacking’s gamma distribution for axon diameter, and the assumption was made that most vessels would fall within a similar size range, as white matter vessel diameter is hardly reported in the literature. Only vessels within the specified size bin were selected to meet the CBV requirement. The default vessel size for other parameter combinations was a random selection across this range until the CBV requirements were met for that simulation. The BOLDsωimsuite package was modified to accommodate the selection of vascular ranges from the input AxonPacking file.

### Axon Spacing

Simulations of interstitial fraction (ISF) were implemented as a change in axon spacing. AxonPacking was used to generate two additional packing distributions with 0.5 μm and 1 μm spacing between axons. This range captures the variability in extracellular space observed in white matter ^27,28^. The default spacing was 0 μm.

### Simulation Methodology

For all simulations, the default parameter values were used as described above unless otherwise specified. Vessels and axons were defined as impermeable ^32^. The apparent diffusion coefficient (ADC) of axons was set to 0.0005 and the ADC of the veins was set to 0.001. Fibre susceptibility was set to −0.0015 ppm (cgs). This value is approximately one order of magnitude smaller than the susceptibility of isolated myelin membranes reported in the range −0.008 to −0.015 ppm (cgs) ^6,35^. The reduction arises because the simulated cylinder represents the full myelinated axon unit rather than the myelin bilayer in isolation. Vessels were either (1) oriented parallel to the fibres (i.e. “parallel”), (2) oriented parallel to the WM fibres 50% of the time and randomly the other 50% of the time (i.e. “mixed”), or (3) entirely randomly oriented (i.e. “random”).

We simulate a spin-echo (SE) sequence with TE=70 ms (Fig. 1c). Fig. 3 shows the simulated SE signal time course as a function of fibre orientation. The orientation dependence of the free-induction decay (FID) sampled at TE=35ms, as a representation of the gradient-echo BOLD signal, is shown in Fig 2a for the oxygenation values tested as an example.

**Figure 2.**
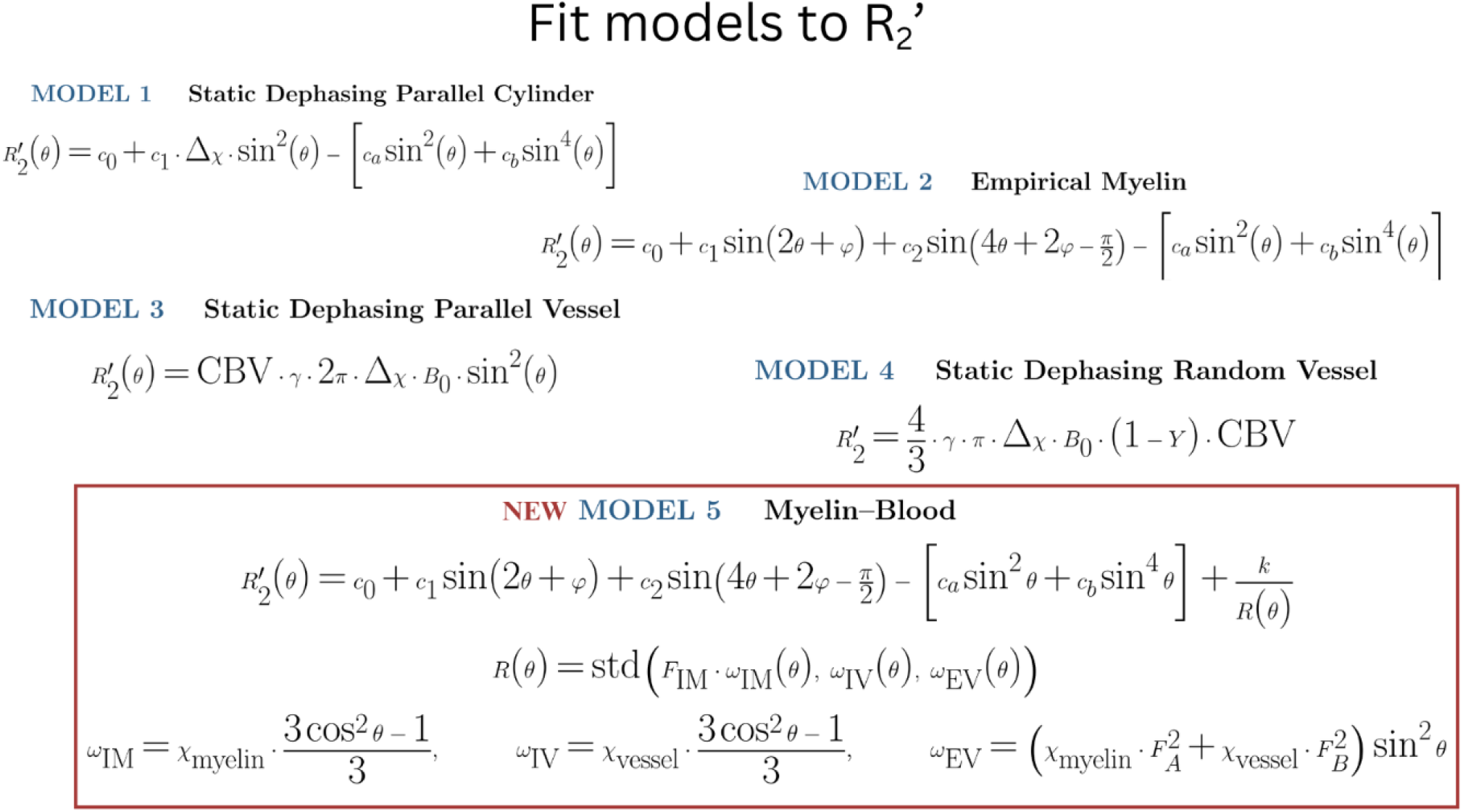
The five models used to fit the R_2_’ data derived from the simulation, with models 1-4 being models reported by previous studies. Models 1 and 2 are for the orientation dependence of R_2_* that were modified to fit R_2_’ by subtracting a fitted term for R_2_. Models 3 and 4 are models developed for R_2_’ specifically. Model 5 is a model developed from Model 2 to include a vascular compartment in an attempt to better fit the R_2_’. The symbols are defined under “Models evaluated”.

**Figure 3.**
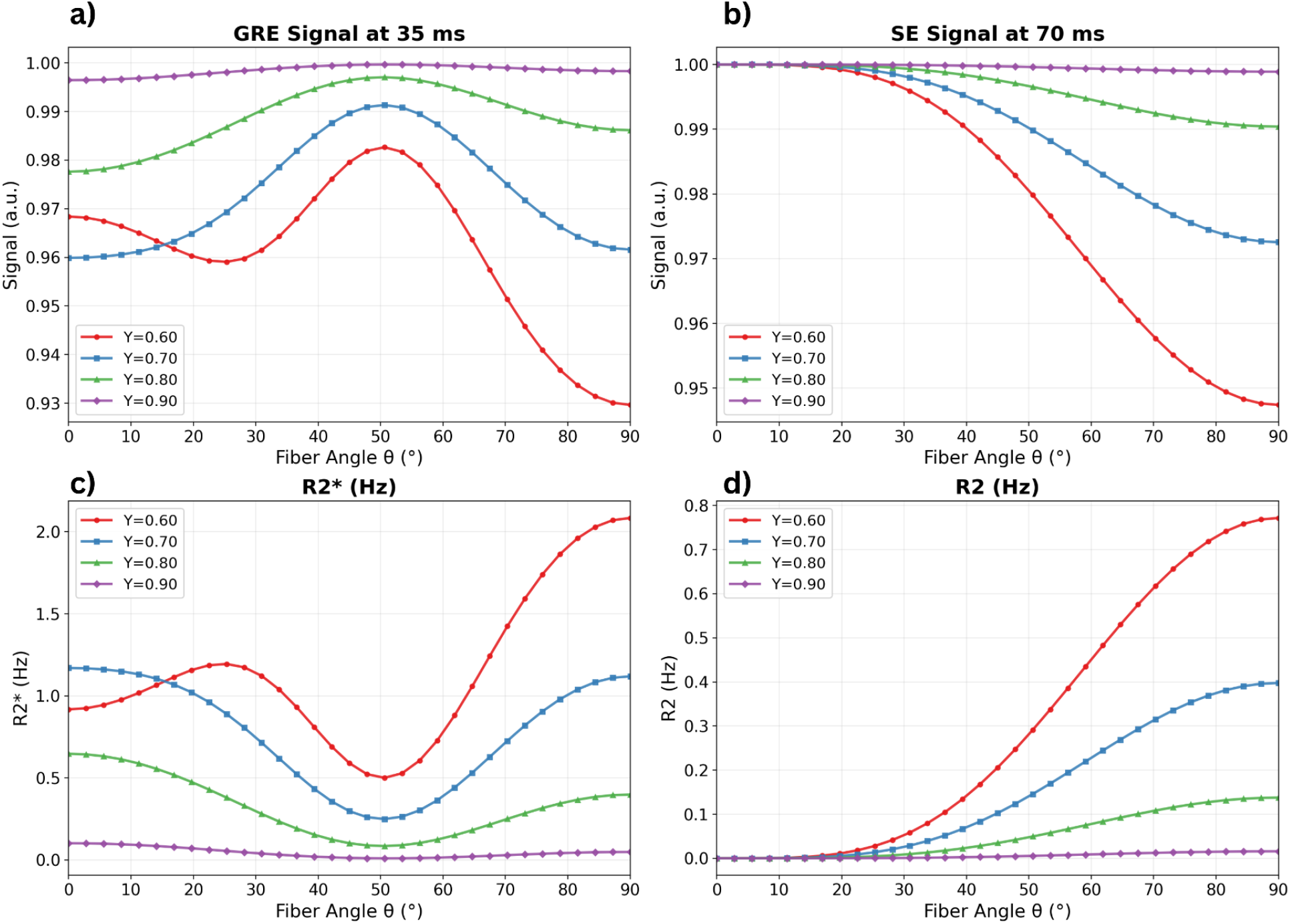
Simulations of the T_2_*- and T_2_-weighted signal as a function of fibre orientation θ. (0° to 90°) for parallel vessel geometry across four blood oxygenation levels (Y = 0.60–0.90). (a) Gradient-echo (GRE) signal at TE = 35 ms. (b) Spin-echo (SE) signal at TE = 70 ms. (c) R_2_* derived from the GRE signal. (d) R_2_ derived from the SE signal. All simulations assumed CBV = 0.02, the medium vessel diameter range of 3.11-5.42 μm, and 0.5 µm axon spacing (38.7% ISF). Curves are coloured by blood-oxygenation level (red: Y = 0.60, blue: Y = 0.70, green: Y = 0.80, purple: Y = 0.90). Both signal and relaxation rates exhibit strong orientation dependence, with a pronounced minimum near θ ≈ 54.7° (the magic angle) visible in R_2_*.

R_2_* and R_2_ were extracted from the FID and SE portions of the signal. R_2_* was calculated using the following formula:

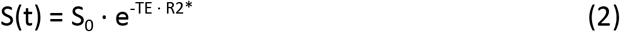

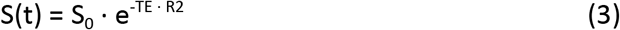

where the TE was 35 ms for GE signal. The same formulation was used for estimating R_2_, where the TE was 70 ms (Fig 2). R_2_’ was calculated as R_2_* − R_2_.

### Models evaluated

We investigated four previously established models to see how well they capture the orientation dependence of R_2_ and R_2_’ in the presence of blood (Fig. 2). In order to represent R_2_’, R_2_ needed to be modeled for removal from R_2_*. Following ^36,37^ and ^15^, who demonstrated that a combination of sin^2^θ and sin^4^θ terms provides the best description of R_2_ orientation dependence in white matter, we adopted this model to describe R_2_. The orientation dependence of R_2_ was therefore modelled as:

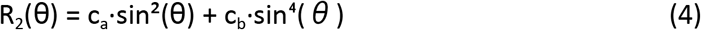

where c_a_ and c_b_ are free-fitted non-negative coefficients (ca, cb ≥ 0). This R_2_ term was subtracted from each of our R_2_* models to isolate the R_2_’ component.

The first model was based on the Static Dephasing Parallel Cylinder model of R_2_* ^3^:

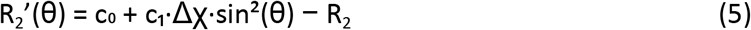

where c_0_, c_1_, are fitted constants and Δχ is the susceptibility difference between oxygenated and deoxygenated blood calculated from Eq. (1) for each oxygenation level. This model has been used to characterize the orientation dependence of myelinated fibres.

The second model was the Empirical Myelin Model ^12^, determined from fixed WM tissue at 7 T:

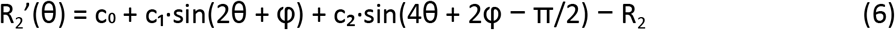

where c_0_, c_1_, c_2_ and φ are constant fitted coefficients. Due to the lack of blood in the post-mortem tissue, this model mainly characterizes the orientation dependence of R_2_* in myelinated fibres.

The third model was the Static Dephasing Parallel Vessel model^3^, as previous work has demonstrated the preference of WM vessels along the fibre direction ^21–23^,

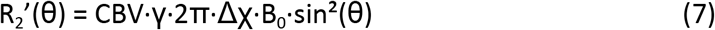

where B_0_ is the field strength, CBV is the cerebral blood volume fraction, and γ is the gyromagnetic ratio 267.513 · 10^6^ rad/s/T. This model was proposed by Yablonskiy and Haacke ^3^ to characterize the R_2_’ in a parallel set of cylinders and is used here to model parallel vessels.

The fourth model used was the static-dephasing random vessel model ^3^, which has been used in the literature to characterize in vivo WM R_2_’ ^38^,

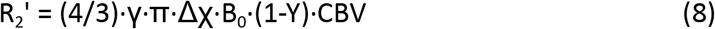

where Y is the blood oxygenation fraction. This model assumes blood vessels are randomly oriented cylinders ^3,38^.

Model fitting employed nonlinear least squares optimization with appropriate initial parameter estimates (Table 1) and was performed in Python. We assessed fit quality using R^2^ and Root Mean Square Error (RMSE). We computed how RMSE varied with each parameter to understand how well each model predicted the R_2_* or R_2_’. Additionally, the absolute percent difference between the simulated R_2_’ and each fitted model was computed and plotted in Fig. 5.

**Table 1.** Initial parameter estimates for model fitting.

| Parameter | Initial value | Models |
| --- | --- | --- |
| *Free parameters (fitted by nonlinear least squares)* |  |  |
| $C_0$ | $\mu$ (mean $R_2'$ of dataset) | 1, 2, 5 |
| $C_1$ | 1.0 | 1, 2, 5 |
| $C_2$ | 0.5 | 2, 5 |
| $\varphi$ | 0.0 | 2, 5 |
| $C_a$ | 0.1 | 1, 2, 5 |
| $C_b$ | 0.1 | 1, 2, 5 |
| $k$ | 1.0 | 5 |
| $F_A$ | 10.0 | 5 |
| $F_B$ | 2.0 | 5 |

### Developing a WM R2’ model that includes the effect of blood vessels

Previous models have derived R_2_’ as orientation-independent ^3^, with a sin^2^θ dependence ^3^ or fit R_2_’ by approximating more complex R_2_* models ^14,38^. Although Models 1,3 and 4 incorporate vascular susceptibility, none simultaneously capture the orientation dependence attributed to myelin microstructure as well as a vascular compartment in one model. This motivated the development of a hybrid model that extends the empirical myelin description ^12^ by adding a vascular dephasing term. The myelin-blood model extends the Empirical Myelin Model by adding a term that captures the dephasing arising from frequency offsets across the intramyelin, intravascular and extravascular water populations. The external field contribution follows the infinite-cylinder formulation ^1,3^, in which the extravascular frequency shift scales as sin^2^θ. Rather than adding a direct sin^2^θ term, however, the model captures the net effect of these compartmental field components through the inverse of their inter-compartmental standard deviation, 1/R(θ).

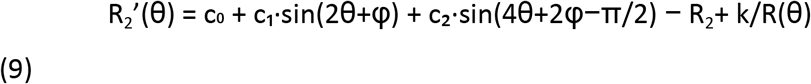

where c_0_, c_1_, c_2_ and φ are the empirical myelin harmonic coefficients, ca and cb (≥ 0) are the non-negative R_2_ subtraction coefficients from the sin^2^θ + sin^4^θ model, and k is a free scalar applied to 1/R(θ). The term 1/R(θ) represents the inverse of the inter-compartmental standard deviation of weighted frequency shifts across the intramyelin (IM), intravascular (IV) and extravascular (EV) water populations:

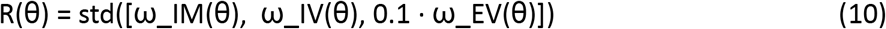

The interior cylinder field shifts are ω_IM = χ_myelin · (3cos^2^θ − 1)/3 and ω_IV = χ_vessel · (3cos^2^θ − 1)/3, where χ_myelin = −0.12 and χ_vessel = +0.24 were held fixed. The extravascular field shift is ω_EV = (χ_myelin · F_A^2^ + χ_vessel · F_B^2^) · sin^2^θ, consistent with the external cylinder field formulation of Ogawa et al. (1993). The standard deviation scaling factors F_A and F_B are free parameters which scale the myelin and vessel contributions to the extravascular field. The extravascular term is additionally down-weighted by a factor of 0.1, reflecting the empirically smaller contribution of the spatially distributed extravascular compartment to the inter-compartmental frequency spread.

## Results

Simulations revealed substantial orientation-dependent effects on both R_2_* and R_2_ across all physiological and microstructural parameter ranges tested, as exemplified in Fig. 4. The detailed results will be discussed in the following sections.

**Figure 4.**
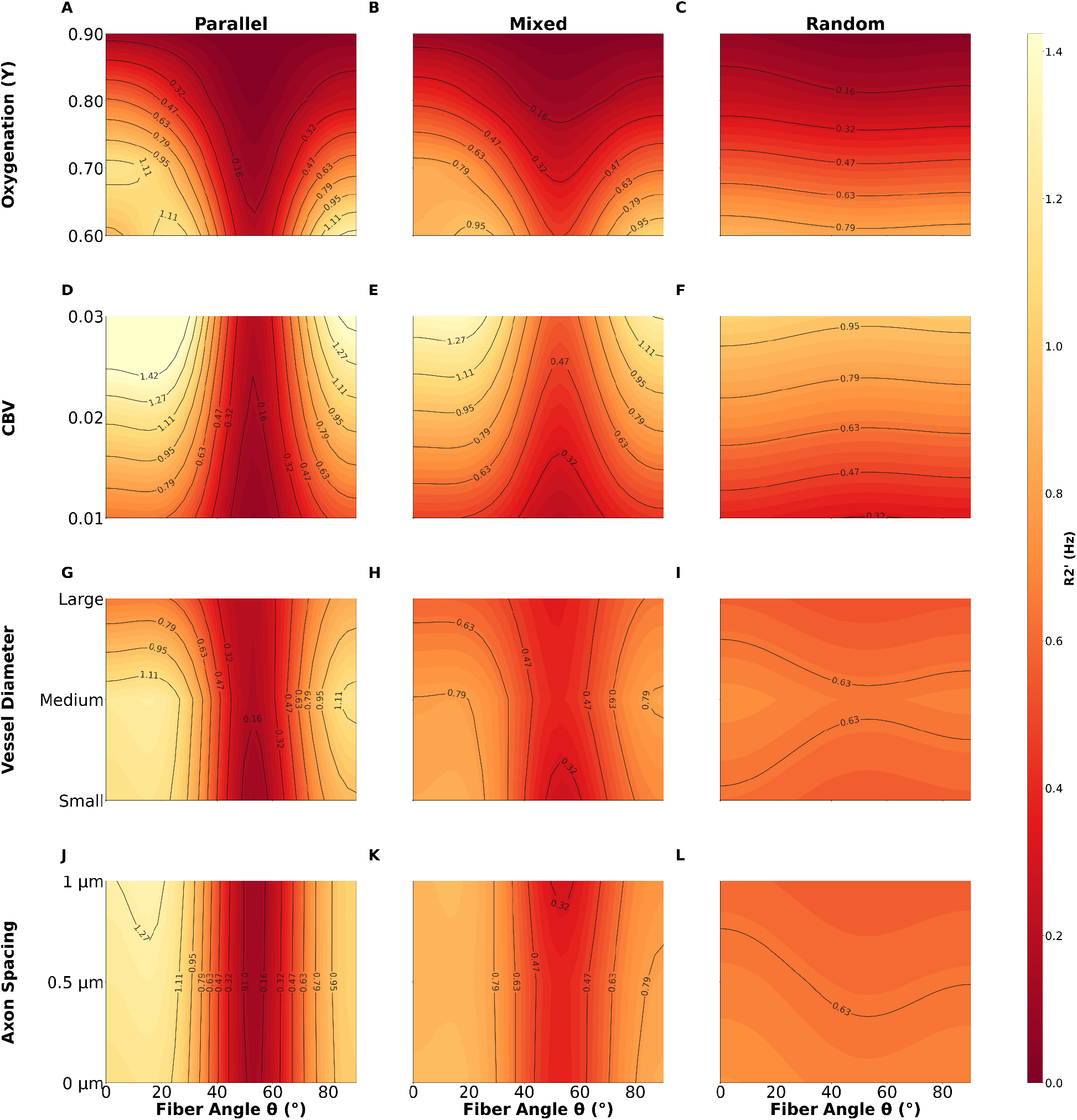
Contour maps of R_2_’ for tested parameter values over range of voxel orientations. The columns from left to right correspond to the parallel, mixed and random vessel geometry, respectively. (a-c) Plots for the blood oxygenation simulations; (d-f) plots for CBV simulations; (g-i) plots for vessel diameter range simulations; (j-l) plots for axon spacing (ISF) simulations. Default parameter values are: Y = 0.7, CBV = 0.02, vessel diameter randomly selected from within the range of 0.80-7.72 μm to meet CBV requirements, with an axon spacing of 0 μm (30.2% ISF).

**Figure 5.**
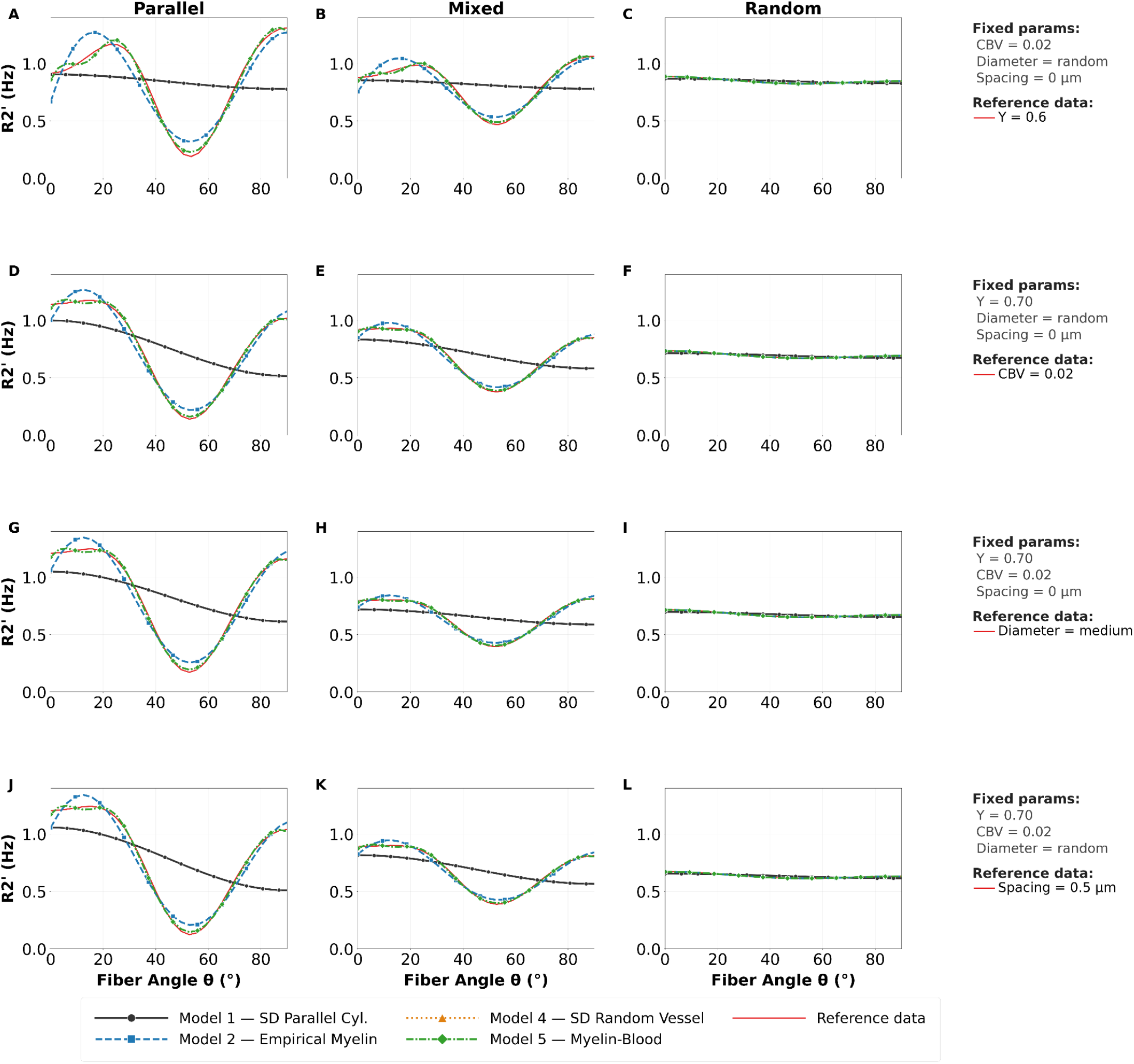
Model fits to sample data from each tested parameter. Four models are shown fitted here, Models 1,2,4 and 5. Model 3 (Static Dephasing Parallel Vessel model) was excluded due to its much larger range and the fit is visible in the supporting information figure S1. Each row shows one parameter sweep across three vessel geometry configurations (parallel, mixed and random). In each panel the red line is the simulated R_2_’ data at the reference condition and the black lines show the four models that fit that reference condition. (A-C) show the model fits to the oxygenation reference condition Y = 0.6. (D-F) show the model fits to the CBV reference condition CBV = 0.02, (G-I) show the fits to the vessel diameter reference condition where the diameter was set to the medium range 3.11-5.42 μm. (J-L) shows the fits to the axon spacing reference condition of 0.5 μm spacing (38.7% ISF). Fixed parameters for each set of simulations are listed in the legend.

### Effect of fibre Orientation

White matter SE and GRE simulated signals exhibited strong dependence on fibre orientation relative to the main magnetic field (Fig. 3). As illustrated for parallel vessel geometries in Figure 2, R_2_ consistently peaked at θ ≈ π/2 (perpendicular orientation). R_2_* exhibited maxima at θ ≈ π/6 and a larger peak at θ ≈ π/2, with a pronounced minima at 53.4°. For random vessel orientations, R_2_’ amplitude was substantially reduced and nearly flat across angles, in contrast to the larger changes seen in parallel and mixed vessel geometries.

### Effect of blood oxygenation

Increasing blood oxygenation (Y) from 0.6 to 0.9 produced decreases in R_2_’ ranging from 0.099 Hz (8.1% change, random vessel orientation) to 1.024 Hz (91.2% change, parallel vessel orientation). The contour plots (Fig. 4a-c) show how R_2_’ changes with Y across fibre angle θ. In parallel vessels, the peak R_2_’ shifts with Y (Fig. 4a). At Y = 0.6 the maximum occurs at θ = 90°, while for Y ≥ 0.7 the maximum shifts to θ = 0°. This shift in peak orientation is visible in the contour plot as a shift in the intensity along the axis as Y increases. R_2_’ also decreased by 1.024 Hz (91.2%) from Y = 0.6 to Y = 0.9 for the parallel vessel orientation. The mixed vessel orientation shows a similar shift in peak R_2_’ along Y, along with a similarly large decrease of 0.513 Hz (86.2%) from low to high Y values. In contrast, the random vessel orientation data showed no shift, and exhibited a near-constant R_2_’ of ~0.065 Hz across all oxygenation levels (8.1% change).

A consistent feature across all vessel orientation distributions is a band of low R_2_’ values centred near θ = 53.4°, which persists across all Y values. The contour lines are densely spaced at low Y and widely spaced for higher values of Y, reflecting the steep reduction in R_2_’ as oxygenation increases.

### Effect of CBV

Variations in CBV (1-3%) produced strong effects on R_2_’ amplitude for parallel and mixed vessel geometries (changes in peak-to-trough range of 1.017 Hz, 172.2% in parallel, 0.548 Hz, 201.2% in mixed). The contour plots (Fig. 4d-f) show a systematic upward shift in R_2_’ intensity across all angles as CBV increases. In parallel vessels, the peak R_2_’ increases from 0.659 Hz at CBV = 0.01 to 1.794 Hz at CBV = 0.03, with the peak consistently located near θ = 15° across all CBV values. Mixed vessels show a similar pattern as parallel vessels, but for random vessel orientations, the contour lines show that R_2_’ varies minimally with changes in θ but tends to be higher with increasing CBV.

The peak-to-trough range in parallel vessels increased by 1.017 Hz (172.2%) from CBV = 0.01 to CBV = 0.03, and similarly for mixed vessels (0.548 Hz, 201.2%), indicating that CBV strongly scales the R_2_’. In contrast, random vessel orientations showed a smaller amplitude change (0.006 Hz, 9.5%).

### Effect of Vessel Diameter

Vessel diameter ranges produced peak-to-trough range changes of 0.386 Hz (36.2%) in parallel and 0.289 Hz (49.2%) in mixed vessel geometries, with negligible effect in random vessel geometries (0.002 Hz, 2.7%). The contour plots (Fig. 4g-i) reveal a notable change in the location of the R_2_’ peak with vessel size for parallel and mixed orientations. For small and medium vessels, the peak occurs near θ = 15.5° for parallel and mixed vessel orientations. For large vessels, the peak shifts to near θ = 90° in both parallel and mixed orientations. In random vessel orientations, the contours and range of values have a smaller range and the peak stays near θ = 0° across all vessel sizes.

Comparing small, medium and large vessel diameters, parallel vessels showed an amplitude decrease of 0.386 Hz (36.2%) from small to large. Mixed vessel orientations exhibited a decrease of 0.289 Hz (49.2%) in amplitude. Random vessel orientations showed essentially no change (0.002 Hz, 2.7%). The position of the angular minima remained consistent at θ ≈ 53° for all three vessel diameter ranges.

### Effect of Axon Spacing

Axon spacing (inter-axonal distance) produced distinct effects across vessel geometries, with parallel and mixed orientations showing increased R_2_’ with greater spacing, and random orientations showing a downward shift in R_2_’ with increased spacing within the simulated voxel.

For parallel vessel orientations, the contour plot (Fig. 4j) shows that both the peak and trough increase with increasing axon spacing which produces an overall peak-to-trough range increase of 13.9%. In mixed vessels, the peak is largely stable, while the trough decreases as spacing increases which again results in a 12.8% increase in amplitude. For random vessel orientations the effect is different: R_2_’ decreases as axon spacing increases from a mean of 0.694 Hz at 0 μm (30.2% ISF) to 0.567 Hz at 1 μm (45.7% ISF).

### Fit to Analytic Models

Model fitting results are organized by model class: vascular models (Models 3 and 4), the myelin-based models (Models 1 and 2), and the novel myelin-blood model (Model 5). Model fit coefficients are averaged across tested values for each parameter and shown for all models tested in supporting information Tables S1-S4.

### Vascular models

The static dephasing random vessel model (Model 4) fit the simulated R_2_’ data poorly across all parameter combinations (mean R^2^ −2158.46, mean RMSE = 1.513 Hz), predicting R_2_’ values 2-3 times larger than the simulated data (Fig. 5). The Static Dephasing Parallel Vessel model (Model 3) performed similarly poorly (mean R^2^ = −1012.42, mean RMSE 1.081 Hz), with the same pattern of overestimation. Although Model 3 captures angular dependence as sin^2^θ, the fixed coefficients substantially overestimate field perturbation magnitudes in both cases.

### Myelin-based models

The Static Dephasing Parallel Cylinder model (Model 1) performed substantially better than the two vascular models but still fit the data poorly overall, with a mean R^2^ of 0.36 and mean RMSE of 0.14 Hz across all conditions. Performance was consistently worse for parallel and mixed vessel orientations compared to random vessel orientations.

The Empirical Myelin Model (Model 2) demonstrated substantially better fits, with a mean R^2^ of 0.981 and mean RMSE of 0.027 Hz across all conditions. Parallel vessel geometries yielded a mean R^2^ of 0.97, mixed orientations gave a mean R^2^ of 0.974 and random vessel orientations showed near perfect fits with near-zero RMSE. Despite these strong metrics, signed error analysis (Fig. 6) reveals a systematic residual structure where Model 2 predominantly overpredicts R_2_’ near the magic angle, where the simulated data approaches zero more steeply than the model. Fit coefficients for the Empirical Myelin Model are shown in Table S5.

**Figure 6.**
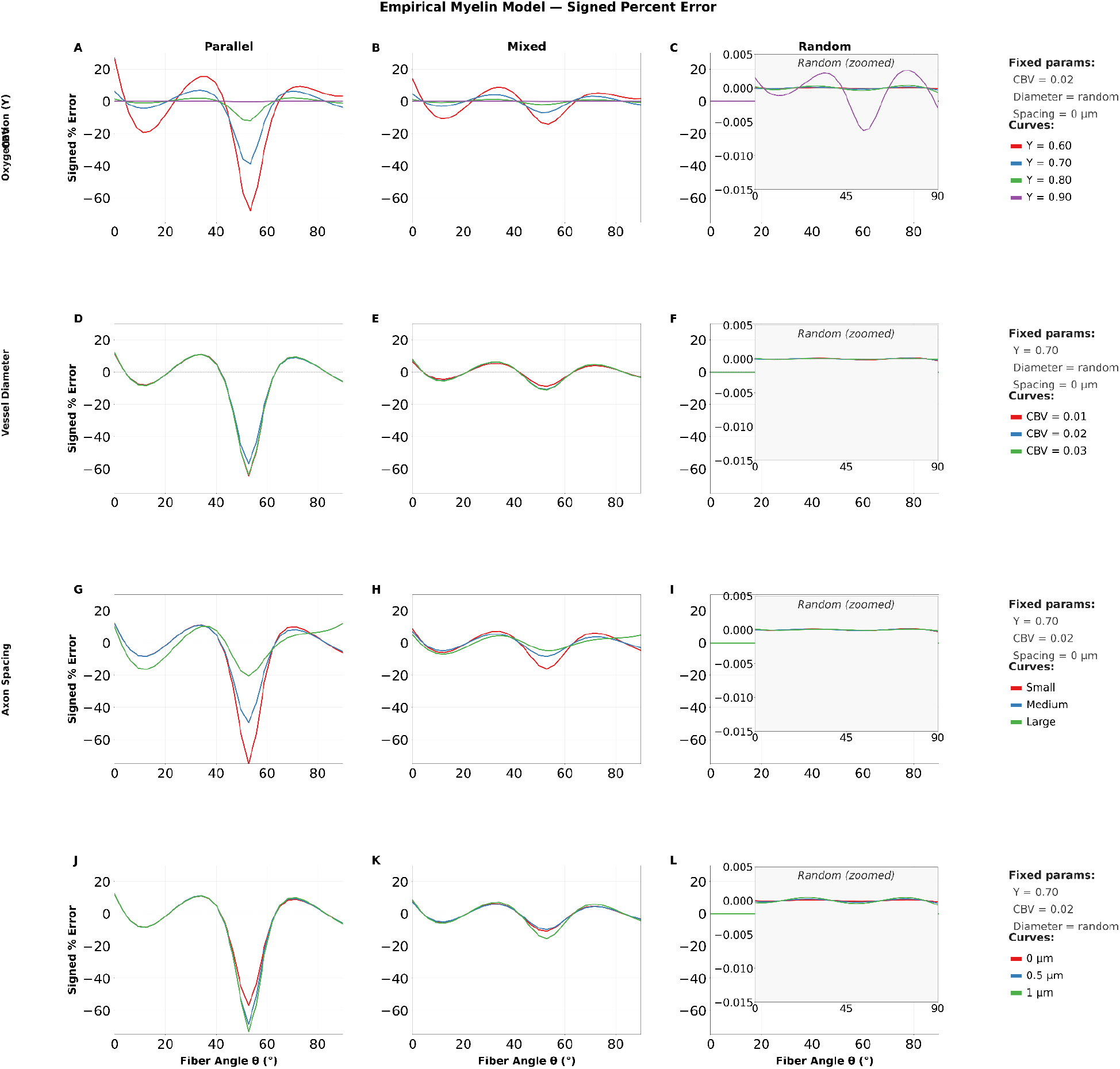
Signed percent error between simulated R_2_’ and the Empirical Myelin model (Model 2) across all parameter sweeps and vessel geometries. Each row shows one parameter sweep across three vessel geometry configurations (parallel, mixed and random). In each panel the error between the simulated R_2_’ and model fits to the data are shown for all parameters simulated. Legends in the last column indicate which line colours represent which data. Random vessel geometry error was much smaller than parallel and mixed vessel geometries, and is shown in an inset window with a smaller scale. (A-C) show the error from the model fits to the oxygenation (D-F) show the error from the model fits to the CBV (G-I) show the error from the fits to the vessel diameter. (J-L) show the fits to the axon spacing (ISF). Signed error is defined as (data-fit)/data x 100, such that negative values indicate the model overpredicts R_2_’ and positive values indicate underprediction. The y-axis is shared across all panels. Unsigned percent error is shown in supporting information figure S2.

### Myelin-Blood Model

The Myelin-Blood model (Model 5) achieved substantially improved fits to the Empirical Myelin Model across all parameters and vessel geometries, with an overall mean R^2^ of 0.999 and mean RMSE of 0.007 Hz. For parallel vessel geometries, Model 5 gave a mean R^2^ of 0.998 (mean RMSE =0.014 Hz, max RMSE = 0.029 Hz), compared with mean R^2^ = 0.970 and max RMSE =0.109 Hz for Model 2, which corresponds to a mean RMSE reduction of 73.7%. Mixed vessel geometries showed a similarly consistent improvement: mean R^2^ = 0.998 (mean RMSE = 0.007 Hz, max RMSE = 0.015 Hz), versus mean R^2^ = 0.974 and max RMSE = 0.055 Hz for Model 2, representing a mean RMSE reduction of 73.8%. For random vessel orientations, both models achieved R^2^ approximately equal to 1, with RMSE approximately equal to 0, which is largely due to the minimal change in the R_2_’. The improvement was most pronounced for parameter combinations which produced the largest vascular susceptibility changes, low oxygenation and large vessel diameters.

Model 5 substantially compressed the signed error range relative to the Empirical Myelin Model, with errors in parallel and mixed geometry panels (Fig. 7) remaining within approximately −20% to 6% across all parameter combinations. The residual signed error structure retains the same qualitative pattern as Fig 5 but at substantially reduced magnitude, indicating that the vessel compartment dephasing term in Model 5 captures a meaningful portion of the residual structure present in the Empirical Myelin Model. Fit coefficients for the Myelin-Blood Model are shown in Table S6.

**Figure 7.**
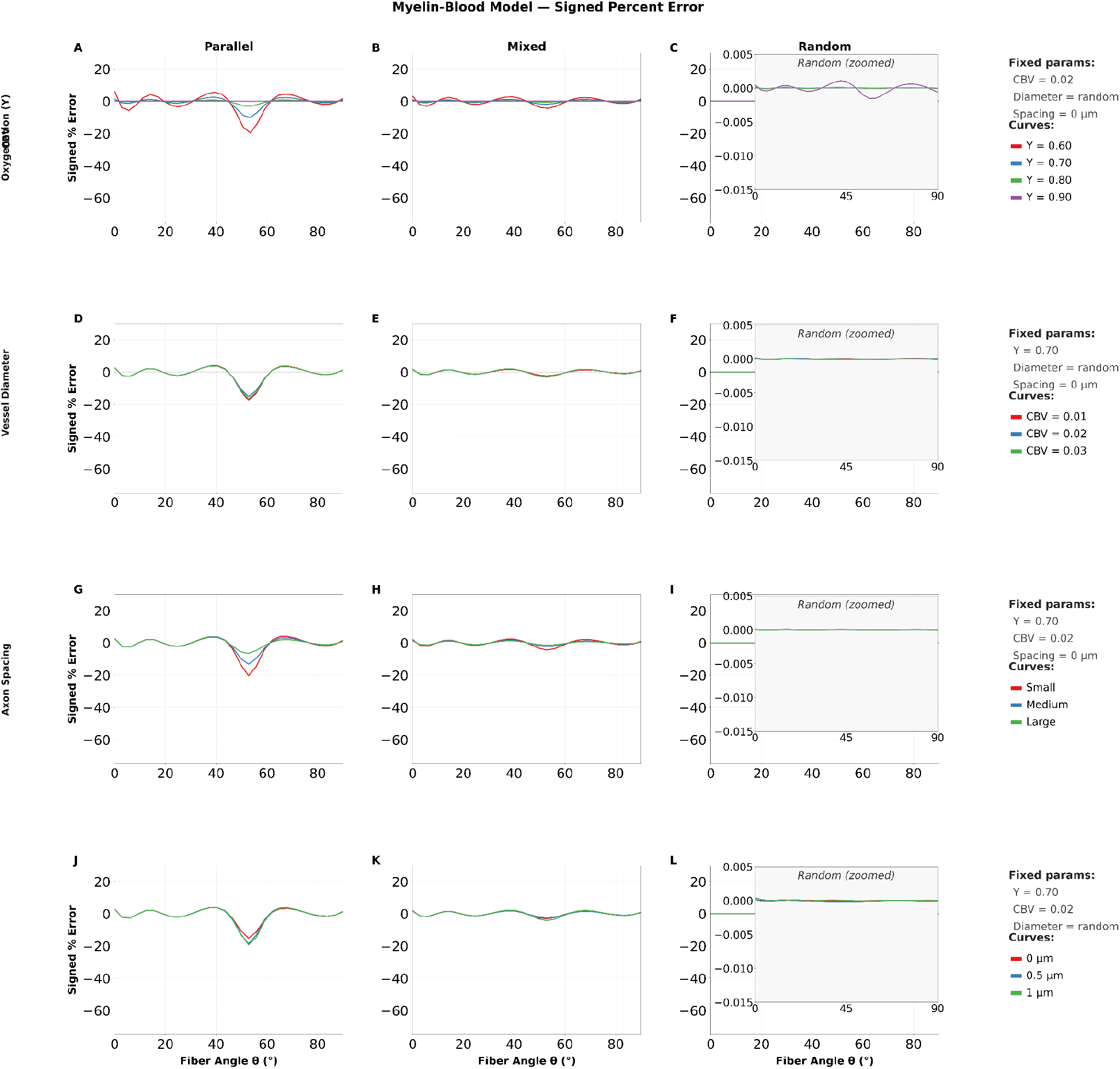
Signed percent error between simulated R_2_’ and the myelin-blood model (Model 5) across all parameter sweeps and vessel geometries. Each row shows one parameter sweep across three vessel geometry configurations (parallel, mixed and random). In each panel the error between the simulated R_2_’ and model fits to the data are shown for all parameters simulated. Legends in the last column indicate which line colours represent which data. Random vessel geometry error was much smaller than parallel and mixed vessel geometries, and is shown in an inset window with a smaller scale. (A-C) show the error from the model fits to the oxygenation (D-F) show the error from the model fits to the CBV (G-I) show the error from the fits to the vessel diameter. (J-L) show the fits to the axon spacing (ISF). The y-axis is shared across all panels. Unsigned percent error is shown in supporting information figure S3.

## Discussion

The reversible transverse relaxation rate R_2_’ plays an important role in MRI investigation of brain physiology. As a quantity sensitive to local magnetic field inhomogeneities, R_2_’ reflects the combined susceptibility environment of the tissue. In particular, for the WM, this encompasses both the paramagnetic effects of deoxyhemoglobin in the vasculature and the diamagnetic susceptibility anisotropy of the myelin sheath ^5,7,8,12^. The present work demonstrates that these two contributions interact in an orientation-dependent manner and that failing to account for this has consequences for two downstream applications: the estimation of the oxygen extraction fraction (OEF) and the interpretation of the WM BOLD signal. Furthermore, we present an analytical model that improves upon the description of WM R_2_’ in current literature and accounts for the presence of both blood and myelin.

### Knowledge gaps in modeling white matter R_2_’

WM fundamentally differs from GM in ways that have not been fully appreciated for R_2_’ specifically. In GM, where vessels are less likely to be strongly oriented with the tissue and myelin is largely absent ^23,26^, R_2_’ can reasonably be treated as reflecting vascular dephasing alone ^3^. In WM however, the ordered architecture of myelinated fibre bundles introduces strong orientation-dependent susceptibility effects. While the orientation dependence of R_2_ and R_2_* in WM has been characterized and attributed primarily to myelin ^6,10,12,14,39^, the equivalent characterization for R_2_’ has been lacking. This has direct consequences for the interpretation of the rapidly emerging area of WM fMRI ^18–20,40^, as well as for quantitative fMRI in the WM ^38^.

These findings have direct consequences for quantitative imaging. OEF estimation via quantitative BOLD (qBOLD) frameworks treats R_2_’ as a purely vascular signal, using it to infer blood oxygenation under the assumption that dephasing is driven by deoxyhemoglobin susceptibility ^3,38,41^. In WM, our results show this assumption is violated: myelin dominates R_2_’ and introduces a large orientation-dependent term that, if unaccounted for, will bias OEF estimates in a direction and magnitude that depends on fibre angle relative to B_0_ ^38^. For WM BOLD the consequences are complementary; because the R_2_’ is orientation dependent and this relies on both the myelin and blood, the sensitivity of R_2_’ to changes in blood oxygenation varies with fibre geometry. Two voxels with the same hemodynamic response may therefore produce different BOLD signal amplitudes depending on the within-voxel vessel orientation relative to B_0_. This is consistent with previous observation of the orientation dependent BOLD signal in WM ^40^ and provides a biophysical basis for the effect. Accurate characterization of WM R_2_’ is thus a prerequisite for reliable interpretation of OEF or the BOLD signal in WM.

The orientation dependence of WM relaxation has been systematically investigated over the past decade, establishing myelin as the primary structural determinant of this effect. Early work demonstrated that both the phase and magnitude of gradient-echo signal in WM vary systematically with fibre orientation relative to B_0_, an effect attributable to the anisotropic magnetic susceptibility of the myelin sheath ^11,12^. Subsequent modeling confirmed myelin as the primary determinant of orientation-dependent R_2_* in WM, with iron playing a secondary role and vascular contributions left unmodelled ^14,42^. This orientation dependence is not limited to gradient-echo contrasts: T_2_-based measures including myelin water fraction also vary with fibre orientation, reflecting the fundamental nature of myelin’s interaction with the magnetic field ^15,36,37^. Notably, however, this body of work has not incorporated an explicit vascular term in the modeling. The present work extends this characterization to R_2_’ and directly addresses the vascular contribution.

### The Role of Diffusion in R_2_’ Simulations

Our choice of the Monte Carlo simulation-based validation framework gets around the challenge of a lack of ground truth, and is the method of choice for capturing the interaction between diffusion and spatially varying dephasing in any WM architecture of our choice ^29–31^. This approach enables the investigation of R_2_’ modeling without the need for knowing WM vascular architecture, which is a concurrent challenge. By simulating proton diffusion explicitly through a voxel containing both myelin cylinders and vascular perturbers with physiologically relevant geometries and hemodynamic parameters, our simulations generate R_2_’ values that reflect the full dephasing physics rather than the static field distribution alone. This provides a more realistic reference signal than analytical models can supply and serves as an in-silico pseudo ground truth against which the analytical and empirical models can be evaluated. The value of Monte Carlo simulation as a tool for model evaluation has been demonstrated previously in the context of quantitative BOLD and vascular MR fingerprinting, where the choice of simulation approach was shown to directly impact parameter estimation accuracy.

### Performance of Existing WM R_2_’ Models and their Limitations

The analytical models used to describe transverse relaxation in the presence of magnetic susceptibility perturbers were developed for a system that does not include diffusion, allowing dephasing to be calculated from the static field distribution alone ^3^. The static dephasing regime provides closed form expressions for R_2_* and R_2_’ that are tractable and have been widely applied, but it explicitly neglects the effect of diffusion on signal formation. As protons diffuse through spatially varying magnetic fields, they sample different field offsets over time, and this averaging modifies the effective dephasing rate in a way that depends on the spatial scale of the field variation relative to the diffusion length. For large vessels, where the field perturbation varies slowly across the extravascular space, diffusion has a limited effect and static dephasing is a reasonable approximation. For small vessels the field varies rapidly on the scale of diffusion length, and protons sample a wider range of field strengths per unit time. In this regime, diffusion substantially accelerates the dephasing rate in ways the static model cannot capture, and analytical approximations that attempt to extend the model to include diffusion are valid only within restricted parameter ranges ^43^.

The models evaluated here span a range of approaches for describing R_2_’ in WM, from purely vascular analytical frameworks to empirically derived myelin-based models, and their relative performance reveals a consistent pattern; the ways in which they do not fit the data reflect the specific assumptions or limitations of each model. The vascular-only models, Parallel Cylinder and Random Vessel (Models 1, 3 and 4), exhibited errors exceeding 50% across most orientations relative to B_0_, with the parallel-vessel model showing particularly large deviations at perpendicular orientations and the Random Vessel model showing errors ranging from 20% at intermediate angles to over 100% at perpendicular orientations. These results highlight that myelin is the primary contributor to WM R_2_’ orientation dependence, and that models attributing all dephasing to vascular susceptibility are missing the primary contributor to the signal. The Static Dephasing Parallel Cylinder model showed systematically higher errors of approximately 10-30% across all angles. This reflects not only the absence of a myelin term but also the limitations of the static dephasing approximation itself; this model assumes the effect of diffusion is negligible, which modifies dephasing around small vessels in ways that static analytical predictions cannot capture ^3,43^. Its uniform error pattern suggests a poor match to the WM signal.

The Empirical Myelin Model (Model 2) performed substantially better than the analytic and vascular-centric models (Models 1, 3 and 4), showing errors typically lower than 20% across most angles capturing the characteristics of the simulated R_2_’ including the trough near the magic angle at θ ≈ 53°. This is consistent with what is known of myelin susceptibility anisotropy in WM transverse relaxation ^5,12,13,39^ and confirms that models derived from myelin microstructure provide a reasonable approximation of WM R_2_’ across a wide parameter space. However, the Empirical Myelin Model exhibited R_2_’ prediction errors >60% near the magic angle, exacerbated by low blood oxygenation values in both parallel and mixed geometries, which are precisely the conditions where vascular susceptibility perturbations are expected to be largest. Critically, the data used for fitting was derived from ex-vivo tissue without a vascular contribution ^12^. The residual errors in these conditions therefore cannot be attributed to noise or model flexibility alone; perhaps requiring a structural term which is missing. Together the performance landscape across these models makes a consistent argument: the biophysics of in vivo WM R_2_’ requires both a myelin term and a vascular term and models that include only one are systematically limited in predictable ways. This motivates the development and evaluation of a model that incorporates both.

### Blood as a Non-Negligible Contributor: The Empirically Informed Myelin-Blood Model

Previous work has suggested that vascular contributions to transverse relaxation are minimal relative to myelin and the performance of the Empirical Myelin Model across much of our parameter space is consistent with myelin microstructure being the dominant determinant of R_2_’ ^6,12,14^. However, dominance is not the same as sufficiency and the residual errors of the myelin only model demonstrate that the vascular contribution, while secondary, is not negligible and cannot be ignored in a complete description of the signal. Notably, prior studies focused on myelin did not include an explicit vascular term, while purely vascular models did not account for myelin, leaving the combined contribution uncharacterized. The incomplete characterization of WM R_2_’ has direct relevance for the rapidly growing field of WM fMRI ^18–20,44^, where orientation-dependent signal changes in R_2_’ complicate the interpretation of task-related responses and resting-state fluctuations in WM tracts. A complete model of WM R_2_’ that accounts for both myelin and vascular contributions is therefore a prerequisite for reliable quantitative WM fMRI.

In this work, we proposed a combined myelin-blood model, which builds on the empirical myelin-dominant model and additionally incorporated a vascular compartment with a term that reflects intra and extravascular boundary dephasing ^1,45^, achieves substantially improved fits across the entire physiological parameter space. Specifically, R_2_’ errors were reduced to below 20% compared to errors of up to 70% for the best-performing existing model (i.e. Empirical Myelin Model), with the most pronounced improvement occurring near the magic angle and at low oxygenation values in parallel and mixed vessel geometries, the same conditions where vascular susceptibility perturbations are largest and where the myelin-only model showed its greatest residual errors. This is mechanistically coherent: the magic angle represents orientations at which myelin’s susceptibility anisotropy is minimized, making the R_2_’ more sensitive to the vascular term. At low oxygenation, the susceptibility difference between intravascular deoxyhemoglobin and surrounding tissue is larger, amplifying the vascular dephasing contribution ^1,34^. The myelin-blood model captures both of these effects because it explicitly represents the vascular compartment as a source of additional orientation-independent dephasing that adds to the myelin driven signal.

The myelin-blood model presented here opens several avenues for advancing quantitative WM imaging. Most directly, it provides a more physically grounded basis for WM-specific qBOLD, in which R_2_’ is used to estimate OEF: incorporating the myelin-driven orientation-dependent baseline into the signal model would allow the vascular component to be isolated more accurately, reducing the fibre-angle-dependent bias that currently limits qBOLD in WM. Beyond OEF estimation, the model offers a framework for interpreting WM BOLD signals in terms of their underlying microstructural and hemodynamic contributors; this enables, in principle, the separation of orientation-dependent signal changes driven by myelin architecture from those driven by genuine hemodynamic responses, especially as the BOLD signal is normalized by the baseline. This would be particularly valuable for WM fMRI studies in tracts with strong orientation biases relative to B_0_, where the two sources of signal variation are currently confounded. The model could also serve as a forward model for vascular MR fingerprinting approaches extended to WM, where dictionaries incorporating both myelin and vascular parameters could enable simultaneous estimation of microstructural and hemodynamic properties from multi-echo data. Each of these applications will require validation against in-vivo data, and the present simulation framework provides the controlled pseudo-ground truth needed to develop and test such approaches before experimental translation.

The present work has several limitations that should be considered in interpreting these findings. The myelin-blood model incorporates a simplified vascular compartment that does not explicitly account for the orientation dependence of the vascular dephasing term itself. In reality, the susceptibility perturbation from blood vessels varies with vessel angle relative to B_0_, which would introduce an additional orientation-dependent vascular contribution not captured by the current orientation-independent implementation. Second, the Empirical Myelin Model used as the foundation for fitting was derived from ex-vivo tissue, which lacks vascular and life-like extra-axonal properties. This could in turn impact our myelin-blood model. Third, the simulations employ idealized single orientation geometries with uniform vessel alignment, whereas real WM voxels contain crossing fibres in 60-90% of cases ^46^ and vasculature with complex branching topology ^21,23^. The extent to which sub-voxel averaging across fibre populations and vessel orientations modifies the R_2_’ signal and the performance of both models remains to be characterized. These limitations define the roadmap for future development of this work toward experimental validation. In future work we aim to assess the feasibility of fitting these models to in-vivo WM data.

## Supporting Information Captions

**Figure S1. Model 3 (Static Dephasing Parallel Vessel model) plotted against the simulation data for oxygenation (Y)**. Dashed lines represent model fits to the respective simulation data, with colours matched for each model/simulation pair.

**Figure S2. Absolute percent error between the simulated R**_**2**_**’ and the Empirical Myelin Model (Model 2) across all parameter sweeps and vessel geometries**. Vessel geometries separated by column. (A-C) show the % error as a function of θ for the oxygenation values tested (Y = 0.6, 0.7, 0.8, 0.9). (D-F) show the % error as a function of θ for the CBV values tested (CBV = 0.01, 0.02, 0.03). (G-I) show the % error as a function of θ for the vessel diameter ranges tested (Vessel diameter: Small = 0.80-3.11 μm, Medium = 3.11-5.42 μm, Large = 5.42-7.72 μm). (J-L) show the % error as a function of θ for the axon spacing distributions tested (Spacing = 0 (30.2% ISF), 0.5 (38.7% ISF), 1μm (45.7% ISF)). Within each panel curves are coloured by parameter value as indicated in the legend. An inset in each random vessel panel shows the data on a zoomed scale, as the random vessel orientation errors are noticeably small.

**Figure S3. Absolute percent error between the simulated R**_**2**_**’ and the Myelin-Blood Model (Model 5) across all parameter sweeps and vessel geometries**. Vessel geometries separated by column. (A-C) show the % error as a function of θ for the oxygenation values tested (Y = 0.6, 0.7, 0.8, 0.9). (D-F) show the % error as a function of θ for the CBV values tested (CBV = 0.01, 0.02, 0.03). (G-I) show the % error as a function of θ for the vessel diameter ranges tested (Vessel diameter: Small = 0.80-3.11 μm, Medium = 3.11-5.42 μm, Large = 5.42-7.72 μm). (J-L) show the % error as a function of θ for the axon spacing distributions tested (Spacing = 0 (30.2% ISF), 0.5 (38.7% ISF), 1μm (45.7% ISF)). Within each panel curves are coloured by parameter value as indicated in the legend. An inset in each random vessel panel shows the data on a zoomed scale, as the random vessel orientation errors are noticeably small.

**Table S1**. Average R^2^ and RMSE for model fits to R_2_’ data from change in blood oxygenation (Y) simulations

**Table S2**. Average R^2^ and RMSE for model fits to R_2_’ data from change in blood volume (CBV) simulations

**Table S3**. Average R^2^ and RMSE for model fits to R_2_’ data from change in vessel diameter simulations

**Table S4**. Average R^2^ and RMSE for model fits to R_2_’ data for change in ISF simulations

**Table S5**. Empirical Myelin Model (Model 2) — fitted parameter means and standard deviations

**Table S6**. Myelin-Blood Model (Model 5) — fitted parameter means and standard deviations

## Acknowledgements

The authors would like to acknowledge financial support from the Canada Research Chairs Program and the Natural Sciences and Engineering Research Council of Canada (J. Jean Chen) and funding support from the Ontario Graduate Scholarship (Nayana Menon).

## Code Availability Statement

The original BOLDsωimsuite package is available at https://github.com/jacobchausse/BOLDswimsuite. Code modifications can be sent upon request to the corresponding author.

