## Supplementary Figures and Tables for "Orientation Dependence of R_2_’ in the White Matter: Digital Characterization, Modelling and Implications for Studying Brain Physiology"

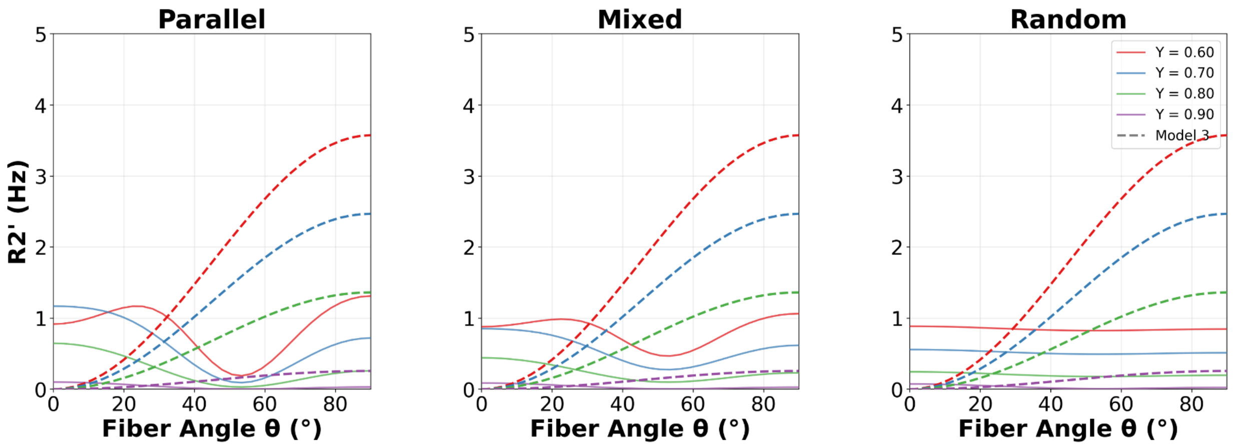


**Figure S1. Model 3 (Static Dephasing Parallel Vessel model) plotted against the simulation data for oxygenation (Y).** Dashed lines represent model fits to the respective simulation data, with colours matched for each model/simulation pair.

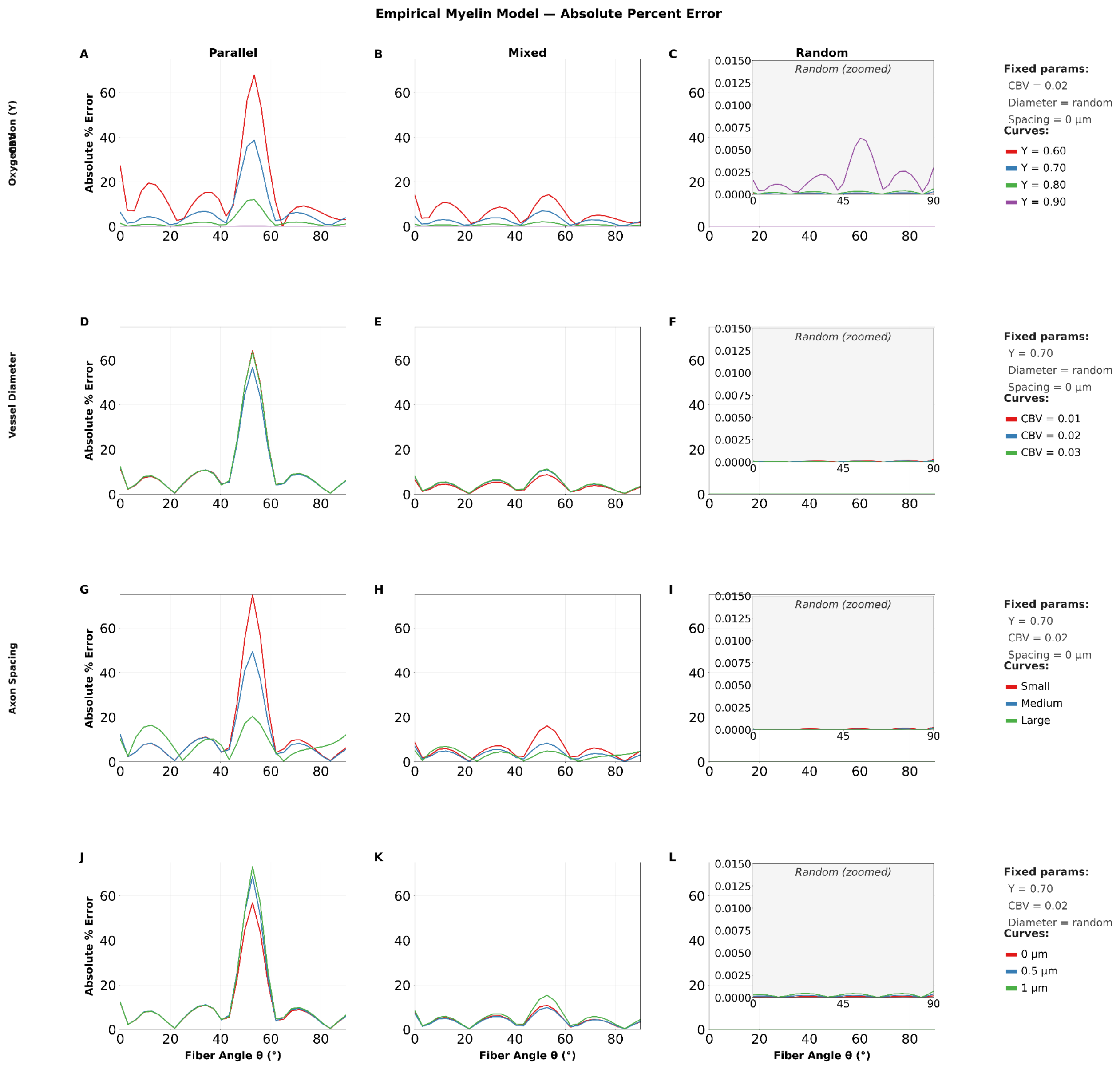


**Figure S2. Absolute percent error between the simulated $R_2$' and the Empirical Myelin Model (Model 2) across all parameter sweeps and vessel geometries.** Vessel geometries separated by column. (A-C) show the % error as a function of θ for the oxygenation values tested (Y = 0.6, 0.7, 0.8, 0.9). (D-F) show the % error as a function of θ for the CBV values tested (CBV = 0.01, 0.02, 0.03). (G-I) show the % error as a function of θ for the vessel diameter ranges tested (Vessel diameter: Small = 0.80-3.11 µm, Medium = 3.11-5.42 µm, Large = 5.42-7.72 µm). (J-L) show the % error as a function of θ for the axon spacing distributions tested (Spacing = 0 (30.2% ISF), 0.5 (38.7% ISF), 1µm (45.7% ISF)). Within each panel curves are coloured by parameter value as indicated in the legend. An inset in each random vessel panel shows the data on a zoomed scale, as the random vessel orientation errors are noticeably small.

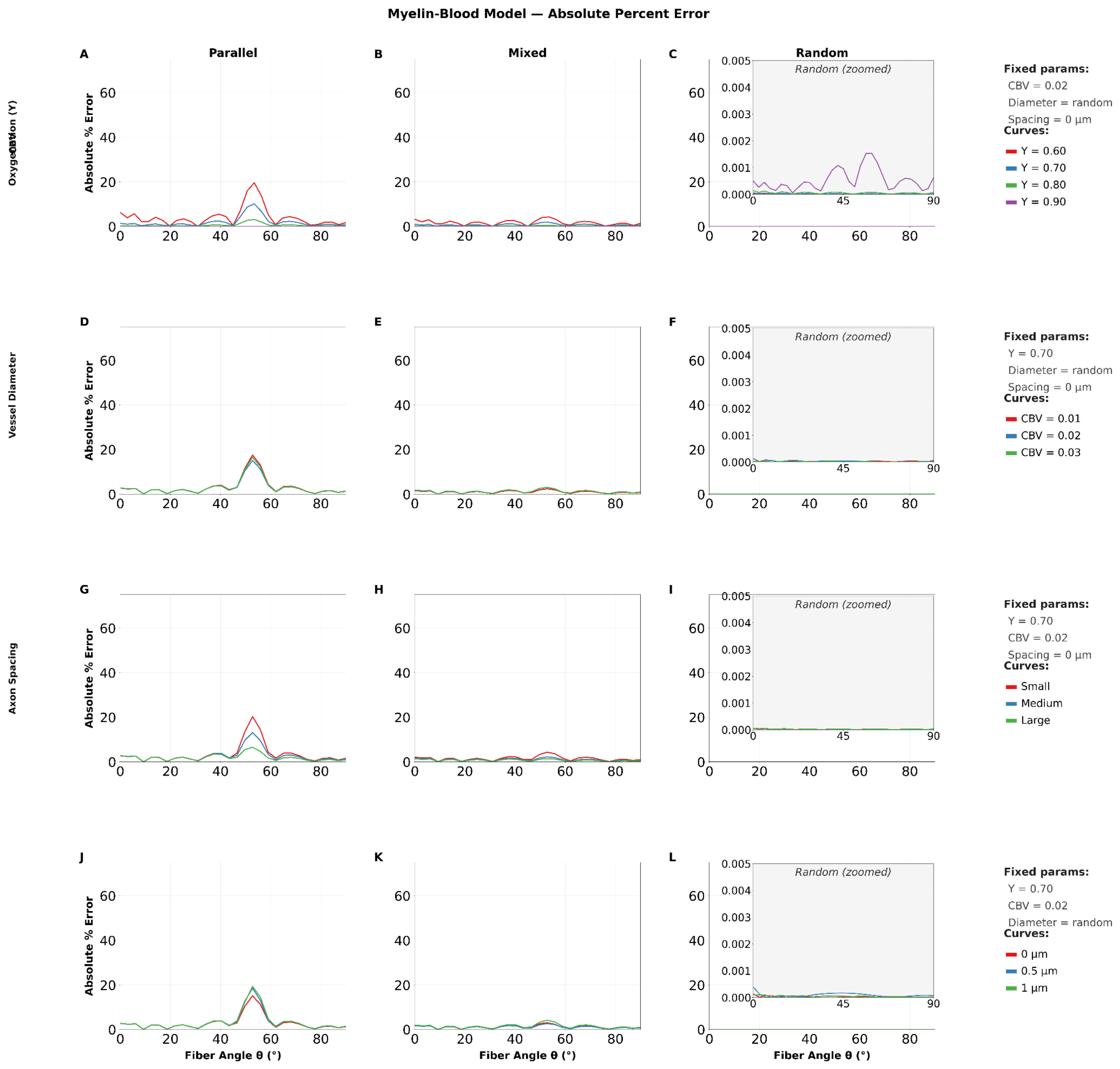


**Figure S3. Absolute percent error between the simulated $R_2'$ and the Myelin-Blood Model (Model 5) across all parameter sweeps and vessel geometries.** Vessel geometries separated by column. (A-C) show the % error as a function of θ for the oxygenation values tested (Y = 0.6, 0.7, 0.8, 0.9). (D-F) show the % error as a function of θ for the CBV values tested (CBV = 0.01, 0.02, 0.03). (G-I) show the % error as a function of θ for the vessel diameter ranges tested (Vessel diameter: Small = 0.80-3.11 µm, Medium = 3.11-5.42 µm, Large = 5.42-7.72 µm). (J-L) show the % error as a function of θ for the axon spacing distributions tested (Spacing = 0 (30.2% ISF), 0.5 (38.7% ISF), 1µm (45.7% ISF)). Within each panel curves are coloured by parameter value as indicated in the legend. An inset in each random vessel panel shows the data on a zoomed scale, as the random vessel orientation errors are noticeably small.

**Table S1.** Average $R^2$ and RMSE for model fits to $R_2'$ data from change in blood oxygenation (Y) simulations

| | **Parallel** | | **Mixed** | | **Random** | |
|---|---|---|---|---|---|---|
| **Model** | $R^2$ | RMSE (Hz) | $R^2$ | RMSE (Hz) | $R^2$ | RMSE (Hz) |
| Model 1 — SD Parallel Cylinder | 0.385 | 0.197 | 0.378 | 0.108 | 0.584 | 0.014 |
| Model 2 — Empirical Myelin | 0.974 | 0.036 | 0.977 | 0.018 | 1.000 | 0.000 |
| Model 3 — SD Parallel Vessel | -16.8 | 0.974 | -45.9 | 0.935 | -2589 | 0.899 |
| Model 4 — SD Random Vessel | -137.2 | 1.422 | -244.8 | 1.437 | -5396 | 1.442 |
| Model 5 — Vascular Hybrid | 0.998 | 0.009 | 0.998 | 0.005 | 1.000 | 0.000 |

**Table S2.** Average $R^2$ and RMSE for model fits to $R_2'$ data from change in blood volume (CBV) simulations

| | **Parallel** | | **Mixed** | | **Random** | |
|---|---|---|---|---|---|---|
| **Model** | $R^2$ | RMSE (Hz) | $R^2$ | RMSE (Hz) | $R^2$ | RMSE (Hz) |
| Model 1 — SD Parallel Cylinder | 0.252 | 0.329 | 0.210 | 0.173 | 0.562 | 0.014 |
| Model 2 — Empirical Myelin | 0.973 | 0.063 | 0.975 | 0.031 | 1.000 | 0.000 |
| Model 3 — SD Parallel Vessel | -9.100 | 1.220 | -32.3 | 1.121 | -3145 | 1.061 |
| Model 4 — SD Random Vessel | -13.9 | 1.492 | -59.5 | 1.513 | -6602 | 1.533 |
| Model 5 — Vascular Hybrid | 0.998 | 0.017 | 0.998 | 0.009 | 1.000 | 0.000 |

**Table S3.** Average $R^2$ and RMSE for model fits to $R_2'$ data from change in vessel diameter simulations

| | Parallel | | Mixed | | Random | |
|---|---|---|---|---|---|---|
| **Model** | $R^2$ | RMSE (Hz) | $R^2$ | RMSE (Hz) | $R^2$ | RMSE (Hz) |
| Model 1 — SD Parallel Cylinder | 0.153 | 0.291 | 0.150 | 0.134 | 0.543 | 0.014 |
| Model 2 — Empirical Myelin | 0.958 | 0.060 | 0.967 | 0.026 | 1.000 | 0.000 |
| Model 3 — SD Parallel Vessel | -16.0 | 1.187 | -78.3 | 1.155 | -2812 | 1.106 |
| Model 4 — SD Random Vessel | -31.4 | 1.551 | -164.3 | 1.629 | -5991 | 1.614 |
| Model 5 — Vascular Hybrid | 0.998 | 0.015 | 0.998 | 0.007 | 1.000 | 0.000 |

**Table S4.** Average $R^2$ and RMSE for model fits to $R_2'$ data for change in ISF simulations

| | Parallel | | Mixed | | Random | |
|---|---|---|---|---|---|---|
| **Model** | $R^2$ | RMSE (Hz) | $R^2$ | RMSE (Hz) | $R^2$ | RMSE (Hz) |
| Model 1 — SD Parallel Cylinder | 0.256 | 0.337 | 0.263 | 0.170 | 0.551 | 0.013 |
| Model 2 — Empirical Myelin | 0.972 | 0.065 | 0.974 | 0.032 | 1.000 | 0.000 |
| Model 3 — SD Parallel Vessel | -8.800 | 1.223 | -32.6 | 1.146 | -3493 | 1.085 |
| Model 4 — SD Random Vessel | -13.5 | 1.481 | -59.9 | 1.540 | -7421 | 1.579 |
| Model 5 — Vascular Hybrid | 0.998 | 0.018 | 0.998 | 0.009 | 1.000 | 0.000 |

**Table S5.** Empirical Myelin Model (Model 2) — fitted parameter means and standard deviations

| | Parallel | | Mixed | | Random | |
|---|---|---|---|---|---|---|
| **Parameter** | **Mean** | **SD** | **Mean** | **SD** | **Mean** | **SD** |
| **Oxygenation (Y)** | | | | | | |
| *c0* | 0.311 | 0.205 | 0.350 | 0.186 | 0.436 | 0.315 |
| *c1* | 0.357 | 0.442 | 0.185 | 0.226 | 0.017 | 0.026 |
| *c2* | -0.447 | 0.350 | -0.238 | 0.180 | 0.004 | 0.024 |
| *φ* | -0.124 | 0.107 | -0.118 | 0.101 | -0.785 | 0.785 |
| *ca* | 0.085 | 0.089 | 0.072 | 0.051 | 0.046 | 0.020 |
| *cb* | 0.138 | 0.164 | 0.052 | 0.057 | 0.034 | 0.037 |
| **CBV** | | | | | | |
| *c0* | 0.544 | 0.199 | 0.581 | 0.213 | 0.719 | 0.261 |
| *c1* | 0.520 | 0.209 | 0.266 | 0.123 | 0.021 | 0.015 |
| *c2* | -0.722 | 0.284 | -0.377 | 0.163 | 0.007 | 0.018 |
| *φ* | -0.264 | 0.002 | -0.242 | 0.006 | -1.047 | 0.740 |
| *ca* | 0.173 | 0.053 | 0.048 | 0.015 | 0.085 | 0.031 |
| *cb* | 0.050 | 0.040 | 0.045 | 0.025 | 0.000 | 0.000 |
| **Vessel Diameter** | | | | | | |
| *c0* | 0.400 | 0.165 | 0.475 | 0.071 | 0.648 | 0.078 |
| *c1* | 0.546 | 0.024 | 0.222 | 0.028 | 0.036 | 0.025 |
| *c2* | -0.672 | 0.103 | -0.296 | 0.061 | 0.025 | 0.005 |

| | | | | | | |
|---|---|---|---|---|---|---|
| *φ* | -0.221 | 0.050 | -0.227 | 0.049 | -1.571 | 0.000 |
| *ca* | 0.019 | 0.027 | 0.037 | 0.052 | 0.071 | 0.056 |
| *cb* | 0.108 | 0.114 | 0.023 | 0.032 | 0.044 | 0.036 |
| **Axon Spacing** | | | | | | |
| *c0* | 0.541 | 0.013 | 0.575 | 0.015 | 0.693 | 0.052 |
| *c1* | 0.540 | 0.026 | 0.263 | 0.021 | 0.047 | 0.012 |
| *c2* | -0.754 | 0.038 | -0.373 | 0.024 | 0.021 | 0.006 |
| *φ* | -0.260 | 0.002 | -0.253 | 0.007 | -1.571 | 0.000 |
| *ca* | 0.096 | 0.068 | 0.082 | 0.022 | 0.102 | 0.017 |
| *cb* | 0.140 | 0.102 | 0.048 | 0.016 | 0.030 | 0.042 |

*Values are mean ± SD of fitted coefficients averaged across parameter values within each sweep. Each row pools all datasets for that parameter (e.g. Y sweep: n=4 per orientation; CBV and Diameter and Spacing sweeps: n=3 per orientation). φ is in radians.*

**Table S6.** Myelin-Blood Model (Model 5) — fitted parameter means and standard deviations

| | Parallel | | Mixed | | Random | |
|---|---|---|---|---|---|---|
| **Parameter** | **Mean** | **SD** | **Mean** | **SD** | **Mean** | **SD** |
| **Oxygenation (Y)** | | | | | | |
| *c0* | 1.385 | 1.274 | 0.920 | 0.812 | 0.423 | 0.310 |
| *c1* | 0.311 | 0.373 | 0.164 | 0.195 | 0.000 | 0.000 |
| *c2* | -0.661 | 0.615 | -0.325 | 0.286 | -0.019 | 0.000 |
| *φ* | -0.036 | 0.027 | -0.034 | 0.027 | -0.000 | 0.000 |
| *ca* | 0.881 | 0.708 | 0.684 | 0.616 | 0.046 | 0.004 |
| *cb* | 0.439 | 0.525 | 0.004 | 0.004 | 0.000 | 0.000 |
| *k* | -0.133 | 0.164 | -0.067 | 0.083 | -0.000 | 0.000 |
| *FA* | 13.396 | 0.834 | 13.033 | 1.156 | 14.650 | 0.150 |
| *FB* | 3.168 | 0.733 | 3.887 | 2.537 | 6.042 | 0.233 |
| **CBV** | | | | | | |
| *c0* | 2.403 | 0.912 | 1.516 | 0.679 | 0.699 | 0.271 |
| *c1* | 0.570 | 0.219 | 0.290 | 0.130 | 0.000 | 0.000 |
| *c2* | -1.197 | 0.479 | -0.605 | 0.217 | -0.019 | 0.000 |
| *φ* | -0.063 | 0.004 | -0.056 | 0.005 | -0.000 | 0.000 |
| *ca* | 1.410 | 0.686 | 0.751 | 0.730 | 0.044 | 0.003 |
| *cb* | 0.923 | 0.718 | 0.394 | 0.284 | 0.000 | 0.000 |
| *k* | -0.241 | 0.093 | -0.119 | 0.053 | -0.000 | 0.000 |

| | | | | | | |
|---|---|---|---|---|---|---|
| *FA* | 12.534 | 0.260 | 13.116 | 0.377 | 12.741 | 1.867 |
| *FB* | 1.704 | 1.114 | 3.302 | 0.798 | 4.442 | 1.910 |
| **Vessel Diameter** | | | | | | |
| *c0* | 2.100 | 0.587 | 1.244 | 0.273 | 0.618 | 0.059 |
| *c1* | 0.492 | 0.125 | 0.220 | 0.069 | 0.000 | 0.000 |
| *c2* | -0.971 | 0.193 | -0.445 | 0.134 | -0.020 | 0.000 |
| *φ* | -0.063 | 0.005 | -0.067 | 0.002 | -0.000 | 0.000 |
| *ca* | 1.647 | 0.843 | 0.749 | 0.221 | 0.043 | 0.002 |
| *cb* | 0.228 | 0.216 | 0.121 | 0.170 | 0.000 | 0.000 |
| *k* | -0.205 | 0.053 | -0.094 | 0.029 | -0.000 | 0.000 |
| *FA* | 12.447 | 0.265 | 13.337 | 0.605 | 14.580 | 0.233 |
| *FB* | 1.309 | 1.120 | 3.881 | 1.187 | 6.291 | 0.016 |
| **Axon Spacing** | | | | | | |
| *c0* | 2.487 | 0.075 | 1.522 | 0.058 | 0.651 | 0.054 |
| *c1* | 0.592 | 0.025 | 0.293 | 0.020 | 0.000 | 0.000 |
| *c2* | -1.220 | 0.170 | -0.613 | 0.042 | -0.017 | 0.002 |
| *φ* | -0.063 | 0.005 | -0.060 | 0.004 | -0.000 | 0.000 |
| *ca* | 1.580 | 1.000 | 0.747 | 0.200 | 0.039 | 0.004 |
| *cb* | 0.838 | 1.099 | 0.457 | 0.149 | 0.000 | 0.000 |
| *k* | -0.249 | 0.010 | -0.122 | 0.008 | -0.000 | 0.000 |
| *FA* | 12.329 | 0.124 | 12.686 | 0.137 | 13.150 | 2.212 |
| *FB* | 0.903 | 0.700 | 2.493 | 0.243 | 4.795 | 2.089 |

*Values are mean ± SD of fitted coefficients averaged across parameter values within each sweep. Each sweep row pools all datasets for that parameter (e.g. Y sweep: n=4 per orientation; CBV and Diameter and Spacing sweeps: n=3 per orientation). φ is in radians.*
